# Balanced nutrient requirements for maize in the Northern Nigerian Savanna: Parameterization and validation of QUEFTS model

**DOI:** 10.1101/609602

**Authors:** Bello M. Shehu, Bassam A. Lawan, Jibrin M. Jibrin, Alpha Y. Kamara, Ibrahim B. Mohammed, Jairos Rurinda, Shamie Zingore, Peter Craufurd, Bernard Vanlauwe, Roel Merckx

## Abstract

Establishing balanced nutrient requirements for maize (*Zea mays* L.) in the Northern Nigerian Savanna is paramount to develop site-specific fertilizer recommendations to increase maize yield, profits of farmers and avoid negative environmental impacts of fertilizer use. The model QUEFTS (QUantitative Evaluation of Fertility of Tropical Soils) was used to estimate balanced nitrogen (N), phosphorus (P) and potassium (K) requirements for maize production in the Northern Nigerian Savanna. Data from on-farm nutrient omission trials conducted in 2015 and 2016 rainy seasons in two agro-ecological zones in the Northern Nigerian Savanna (i.e. Northern Guinea Savanna “NGS” and Sudan Savanna “SS”) were used to parameterize and validate the QUEFTS model. The relations between indigenous soil N, P, and K supply and soil properties were not well described with the QUEFTS default equations and consequently new and better fitting equations were derived. The average fertilizer recovery fractions of N, P and K in the NGS were generally comparable with the QUEFTS default values, but lower recovery fractions of these nutrients were observed in the SS. The parameters of maximum accumulation (*a*) and dilution (*d*) in kg grain per kg nutrient for the QUEFTS model obtained were respectively 35 and 79 for N, 200 and 527 for P and 25 and 117 for K in the NGS zone and 32 and 79 for N, 164 and 528 for P and 24 and 136 for K in the SS zone. The model predicted a linear relationship between grain yield and above-ground nutrient uptake until yield reached about 50 to 60% of the yield potential. When the yield target reached 60% of the potential yield (i.e. 6.0 tonnes per hectare), the model showed above-ground nutrient uptake of 19.4, 3.3 and 23.0 kg N, P, and K, respectively, per one tonne of maize grain in the NGS, and 17.3, 5.3 and 26.2 kg N, P and K, respectively, per one tonne of maize grain in the SS. These results suggest an average NPK ratio in the plant dry matter of about 5.9:1:7.0 for maize in the NGS and 3.3:1:4.9 for maize in the SS. There was a close agreement between observed and parameterized QUEFTS predicted yields across the two agro-ecological zones (R^2^ = 0.70 for the NGS and 0.86 for the SS). We concluded that the QUEFTS model can be used for balanced nutrient requirement estimations and development of site-specific fertilizer recommendations for maize intensification in the Northern Nigerian Savanna.

## 1. Introduction

The average number of individuals facing food insecurity in Nigeria increased from 40.7 million between 2014 to 2016 to 46.1 million between 2015 to 2017 (FAOSTAT, 2018a). Maize (*Zea mays* L.), the most widely grown arable crop (Adesoji et al., 2016) and valuable cereal in Nigeria (FAO, 2016), can play a vital role in achieving food security in the country providing that the current meagre yield of the crop is increased drastically. Grain yield of maize in Nigeria over the last several decades has been hovering at 2 tonnes per hectare (t ha^-1^) (FAOSTAT, 2018b), which is far less than the yield of about 7 t ha^-1^ observed in well-managed field experiments (Fakorede, 2003; Sileshi et al., 2010). One of the plausible reasons for the huge maize yield gap in Nigeria, as in other many countries in Sub-Saharan Africa, is poor soil fertility, the result of inherently low soil nutrient reserves as well as continuous cropping with inadequate nutrient replenishment (Manu et al., 1991; Ekeleme et al., 2014).

The Northern Nigerian Savanna (especially the Northern Guinea Savanna agroecology) is the most suitable zone for maize production in Nigeria due to high incident solar radiation, adequate rainfall, moderate incidences of biotic stresses and natural dryness at the time of harvest. However, soils in the Northern Nigerian Savanna are the major limitation for maize production intensification. They are predominantly sandy Lixisols, Acrisols, and Cambisols with low activity clays (like kaolinite), small organic matter contents and small nutrient reserves, and prone to water and wind erosion (FDALR, 1999; FFD, 2012; Jones and Wild, 1975). Use of Fertilizer in maize production is necessary in this environment to replenish nutrients removed through the harvested product and exported crop residues (a common practice by most farmers in the area). Fertilizer use for maize production in the Northern Nigerian Savanna as the case in other agroecological zones of Nigeria, has been conventionally promoted through blanket recommendations regardless of wide variability in soil, climate and management regimes. The use of blanket fertilizer recommendations, however, is bound to create imbalanced crop nutrition since maize is cultivated in highly heterogeneous fields (Kihara et al., 2016; Shehu et al., 2018). Such imbalances lead to increased nutrient losses and low fertilizer use efficiency (Cassman et al., 2002), which can impede productivity, profitability and sustainability of a farm (Ezui et al., 2016). To reduce the persistent maize yield gaps in the Northern Nigerian Savanna, appropriate fertilizer recommendations need to be developed based on establishing balanced nutrient requirements, for specific yield targets and tailored to account for a specific field and/or soil condition.

A balanced requirement of a given nutrient refers to an amount of the nutrient required to meet a plant’s needs while maximizing the use efficiency of the nutrient (Ezui et al., 2016). When more than one nutrient is needed, for example, nitrogen (N), phosphorus (P) and potassium (K), balanced requirements refer to optimization of use efficiency of these three nutrients and simultaneously resulting in the largest response to their supplies (Ezui et al., 2016). The QUantitative Evaluation of the Fertility of Tropical Soils (QUEFTS) is a practical model that can be used to estimate balanced nutrient requirements for a location and for a target yield level while accounting for the interactions among macronutrients (particularly N, P and K) that affect plant’s physiological efficiencies (Janssen et al., 1990). The original QUEFTS model was developed for maize using data from Suriname and Kenya (Janssen et al., 1990) and it was later improved by Smaling and Janssen (1993) and Sattari et al. (2014). The QUEFTS model has been successfully tested for other crops like rice, wheat, cassava and sweet potato in different regions (Witt et al., 1999; Pathak et al., 2003; Ezui et al., 2016; Lam et al., 2016). Four major steps are involved in QUEFTS modelling (Sattari et al., 2014); (i) potential supply of the available nutrients (N, P and K) is calculated depending on the indigenous soil supply of the nutrient, plus average fertilizer recovery fraction multiplied by the amount of nutrient input. The indigenous soil nutrient supply is estimated by applying relations between soil chemical properties of the 0-20cm soil layer and dry matter uptake of the nutrient in plots where this very nutrient is omitted; (ii) actual uptake of each nutrient is calculated based on the potential supply of that nutrient, considering the potential supply of the other two nutrients; (iii) the establishment of yield ranges as a function of uptake of the nutrients for maximum dilution and accumulation of that nutrient, respectively; and (iv) the yield ranges are combined into pairs, and yield estimated for pairs are averaged to obtain an ultimate yield estimate considering the maximum potential yield of the crop.

The most fickle part of QUEFTS model is the relationship between soil chemical characteristics and the supply of available nutrients described in step 1 (i) above, as many local environmental factors may interfere (Sattari et al., 2014). In the original version of QUEFTS model the soil supply of available nutrients is calculated from soil chemical characteristics using regression equations primarily requiring datasets of soil organic carbon, available P, exchangeable K and pH (Janssen et al., 1990). The applicability and effectiveness of these default QUEFTS indigenous soil nutrient supply equations in different environments other than those which the model was developed is uncertain. Tabi et al. (2008) applied the QUEFTS model in maize to quantify potential supply of soil N and P, utilization efficiency and fertilizer recovery fractions in Northern Nigeria. This study was based on experiments conducted in only 27 farmers’ fields in two villages, limiting their representativeness for the entire maize producing area in the Northern Nigerian Savanna. It follows that it remains necessary to parameterize and validate the QUEFTS model to obtain balanced nutrient requirements for maize production at scale in the Northern Nigerian Savanna to enable effective implementation of site-specific nutrient management (SSNM) practices. The objectives of this study were to: (1) assess the relation between indigenous soil nutrient supply and soil chemical characteristics in the Northern Nigerian Savanna, (2) parametrize standard coefficients of QUEFTS model to determine balanced nutrient requirements for maize in the Northern Nigerian Savanna, and (3) validate the performance of the QUEFTS model in predicting maize grain yield in the Northern Nigerian Savanna.

## 2. Materials and Methods

### 2.1 Site selection, description and experimental design

To generate datasets for this study, on-farm nutrient omission experiments were conducted over two rainy seasons (2015 and 2016) across fourteen study sites in three administrative States of the Northern Nigerian Savanna (Shehu et al., 2018). The three administrative States included Kaduna (with experimental fields in Lere, Kauru, Soba, Ikara, Makarfi, and Giwa local government areas), Katsina (with experimental fields in Funtua, Dandume, Faskari and Bakori local government areas) and Kano (with experimental fields in Tofa, Bunkure, Tudun Wada and Doguwa local government areas) (Fig. 1). The study sites were chosen to cover a broad range of maize growing conditions across the high production potential areas in the Northern Nigerian Savanna and to involve areas where research for development can support extension support programmes engaged in maize value chain initiatives. In overall the study sites fell within two agro-ecological zones (Fig. 1) i.e. the Northern Guinea Savanna (NGS) and Sudan Savanna (SS). The weather conditions of the two agro-ecological zones during the two years of experimentation are summarized in Fig. 2. The total annual rainfall in NGS was 1128 mm in 2015 and 1130 mm in 2016; total annual rainfall in SS was 717 mm in 2015 and 771 mm in 2016. Experimental fields were selected by generating one or two 10 × 10 km grid(s) in each study site (depending on the size of the study site) using ArcGIS software (Environmental System Research Institute, Redlands, CA, USA). Within each of these 10 km × 10 km grid(s), five 1 km × 1 km sub-grids were delineated evenly. In each of the 1 km × 1 km sub-grids, a field for experimentation was randomly selected, considering the willingness of a farmer and availability of land for the trial setup. A total of ninety-five (95) and one hundred and three (103) experimental fields were selected in the 2015 and 2016 rainy seasons, respectively (Fig. 1). At each experimental field, two sets of trials were established side by side; one with hybrid maize (hybrid) and the other one with open-pollinated maize (OPV).

**Fig. 1:**
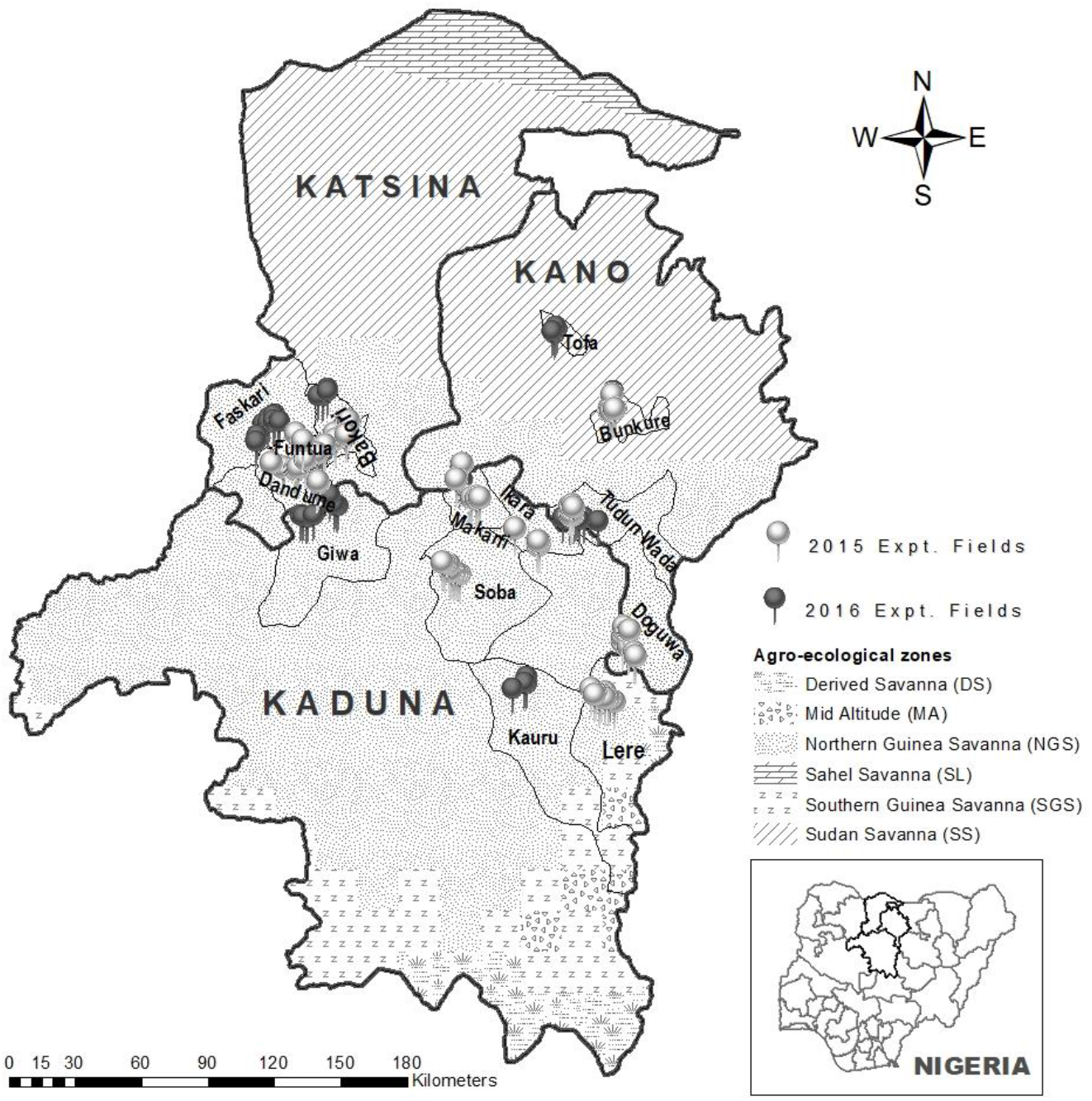
A map of Nigeria showing agroecological zones (AEZ), study sites and experimental fields for on-farm diagnostic nutrient omission trials (NOTs) established in 2015 and 2016 cropping seasons.

**Fig. 2:**
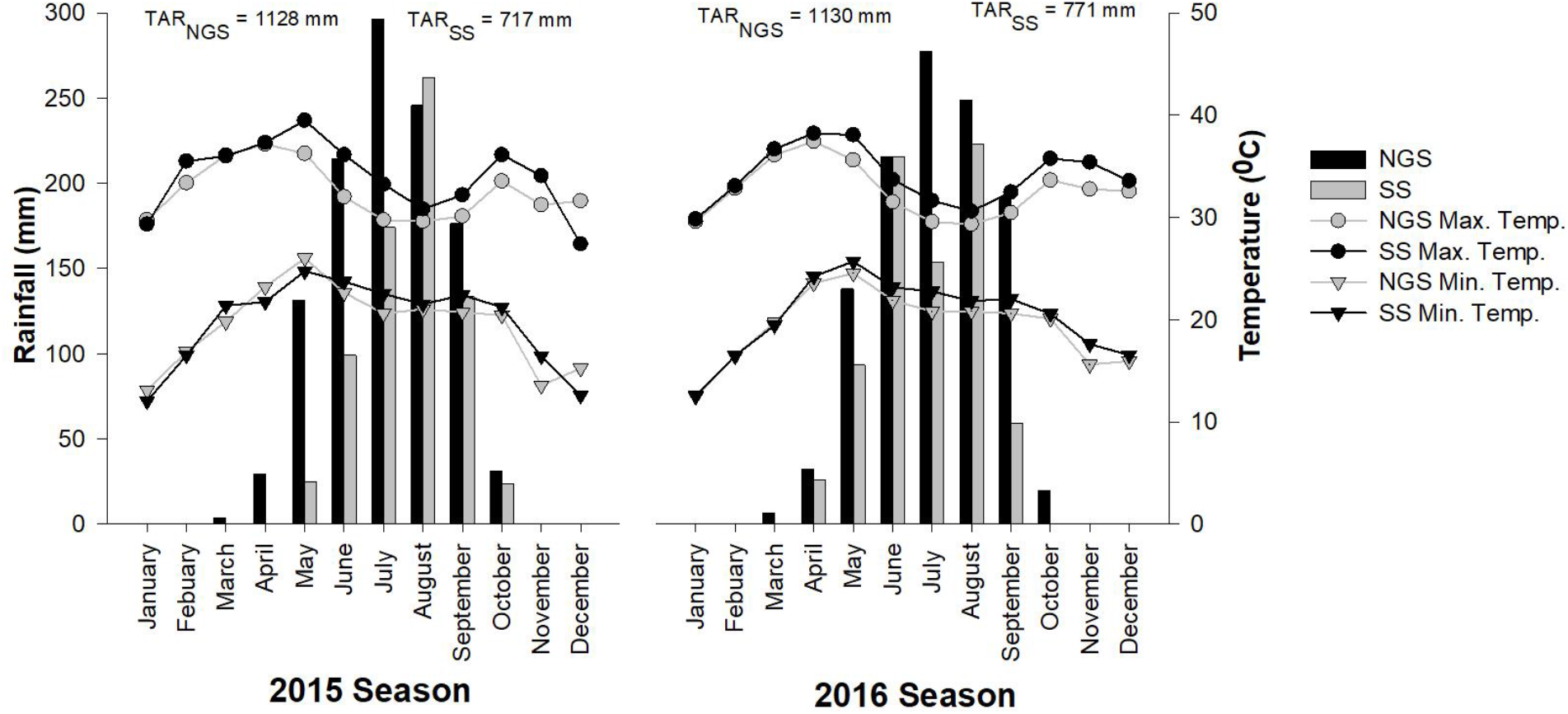
Annual rainfall, daily minimum and maximum temperatures of the two studied agroecological zones recorded in two cropping seasons (2015 and 2016). NGS: Northern Guinea Savanna; SS: Sudan Savanna; TAR_NGS_: total annual rainfall in NGS; TAR_SS_: total annual rainfall in SS; Min.: minimum; Max.: maximum; Temp.: temperature.

The nutrient omission experiments were composed of six nutrient application treatments: (i) control without nutrients applied (control), (ii) N omitted with P and K applied (-N), (iii) P omitted with N and K applied (-P), (iv) K omitted with N and P applied (-K), (v) treatment with all the three nutrients applied (NPK), and (vi) a treatment where secondary macronutrients (S, Ca and Mg) and micronutrients (Zn and B) were applied in addition to the NPK (NPK+). Primary macronutrients were applied at 140 kg N ha^-1^, 50 kg P ha^-1^ and 50 kg K ha^-1^ for the NGS sites; and at 120 kg N ha^-1^, 40 kg P ha^-1^ and 40 kg K ha^-1^ for the SS sites. The secondary macro- and micro-nutrients were applied at 24 kg S ha^-1^, 10 kg Ca ha^-1^, 10 kg Mg ha^-1^,5 kg Zn ha^-1^ and 5 kg B ha^-1^ at all sites. Nitrogen (N) was applied in three equal splits, i.e. at planting (basal application), at 21 and 42 days after emergence (DAE), while all other nutrients were applied at planting. The open-pollinated maize varieties used were IWD C2 SYN F2 (with 105–110 days to maturity) and EVDT W STR (with 90–95 days to maturity) in the NGS and the SS study sites, respectively. The hybrid maize varieties used were OBA SUPER-9 (with 105–110 days to maturity) and OBA SUPER-1 (with 105–118 days to maturity) in all the study sites for 2015 and 2016 seasons, respectively. Treatment plot size was 5 m × 6 m (30 m^2^) with a plant spacing of 0.75 m (inter-row) and 0.25 m (intra-row). Detailed information about the nutrient omission trials is provided by Shehu et al. (2018).

### 2.2. Field and laboratory measurement

Four auger soil samples were collected from 0–20 cm depths from each experimental field during trial establishment before application of fertilizer treatments using a zig-zag random sampling pattern. The four collected samples were thoroughly mixed to have one disturbed composite sample per experimental field and passed through a 2mm sieve for laboratory analysis. Total soil organic carbon (OC_tot_) was assessed using a modified Walkley & Black chromic acid wet chemical oxidation and spectrophotometric method (Heanes, 1984). Total nitrogen (N_tot_), was determined using a micro-Kjeldahl digestion method (Bremner, 1996). Soil pH in water (soil/water ratio of 1:1) was measured using a glass electrode pH meter and the particle size distribution with the hydrometer method (Gee and Or, 2002). Available phosphorus (P_av_), available sulphur (S_av_), exchangeable cations (K, Ca, Mg and Na) and micronutrients (Zn, Fe, Cu, Mn and B) were analysed based on the Mehlich-3 extraction procedure (Mehlich, 1984) preceding inductively coupled plasma optical emission spectroscopy (ICP-OES, Optima 800, Winlab 5.5, PerkinElmer Inc.,Waltham, MA, USA). Exchangeable acidity (H + Al) was determined by extracting soil with 1N KCl and titration of the supernatant with 0.5M NaOH (Anderson and Ingram, 1993). Effective cation exchange capacity (ECEC) was calculated as the sum of exchangeable cations (K, Ca, Mg and Na) and exchangeable acidity (H + Al).

The crop was harvested at physiological maturity in a net plot of 9 m^2^ (i.e. comprising four middle rows of 3 m length of the experimental plot). Plants in the net plot were harvested, and total fresh weights of cobs and stover were recorded. Ten cobs and five stalks of stover were randomly selected as subsamples for nutrient analysis and to account for grain shelling percentage and moisture content after air-drying. The random selection was carried out by first counting the number of cobs or stalks in the net plot and then randomly arranging them in line; the sub samples were then taken at every interval calculated as the total number of cobs or stalks in the net plot over the number of sub samples to be taken. Finally, grain yield was expressed on a dry weight basis at 15.0% moisture content and the stover yield was expressed on an oven dried basis (dried at 60°C). The concentration of total nitrogen in the grain and stover was determined using a micro-Kjeldahl digestion method (Bremner, 1996), while P and K were analysed by digestion with nitric acid (HNO_3_) and concentrations measured with inductively coupled plasma optical emission spectroscopy (ICP-OES, Optima 800, Winlab 5.5, PerkinElmer Inc.,Waltham, MA, USA).

### 2.3. Data Screening and Analysis

Multivariate outliers (n=219) from the experimental data were discarded first at **P<0.05 using Mahalanobis distance in JMP version 13.0 statistical software (SAS Institute Inc., 2017). To understand the characteristics of the screened experimental data (n=1371), analysis of variance was computed using the same JMP 13.0 statistical software. Nutrient application (NA), agro-ecological zone (AEZ) and variety group (VG) were used as main factors. Season was unincluded in the ANOVA because different fields were used between the two seasons of the field experimentation. Mean values with significant differences were compared using Tukey’s HSD (Honestly Significant Difference) test. Finally, the screened experimental data was randomly divided into 80% independent fields for parameterization (n=1090) and the remining 20% (n=281) for validation of the QUEFTS model.

### 2.4. QUEFTS model parameterization and validation

#### 2.4.1 Model parameterization

*Step 1 (assessment of the supply of available nutrients)*: the supply of available nutrients (*S*) in the QUEFTS model is given as a function of indigenous soil nutrient supply plus the nutrient input supply. The nutrient input supply is a function of the quantity of nutrient input added multiplied by the average fertilizer recovery efficiency. The indigenous nutrient supply was developed using a multiple linear regression between soil properties and the uptake of the nutrient in the omitted plots using a stepwise selection procedure. The fertilizer recovery efficiency (*R_i_*) is then calculated as:

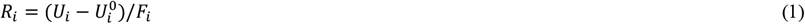

Where *U_i_* = *i*^th^ nutrient in the above ground biomass (kg ha^-1^) in the NPK plot, 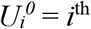 nutrient in the above ground biomass (kg ha^-1^) in the omission plot, *F_i_* = amount of i^th^ nutrient applied (kg ha^-1^).

*Step 2 (relation between the supply of available nutrients and actual uptake)*: The relations between supply of nutrients and actual uptake were calculated using the following conditions and functions (Janssen *et al.,* 1990; Sattari *et al.,* 2014):

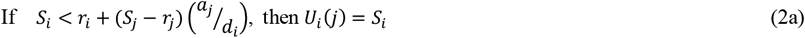

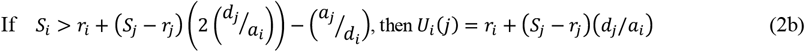

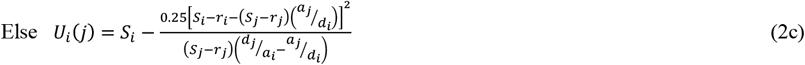

Where *i,j* = *N,P,K, i* ≠ *j*; *U_i_*(*j*) = refers to uptake of *i*^th^ nutrient in relation to *j*, if *i* = N, j may be P or K; *S_i_* = supply of available *i*^th^ nutrient obtained from step 1; *a_i_* = physiological efficiency (*PhE*) or internal efficiency (*IE*) at maximum accumulation of nutrient *i* (kg grain kg^-1^ nutrient *i*); *d_i_* = physiological efficiency (*PhE*) or internal efficiency (*IE*) at maximum dilution of nutrient *i* (kg grain kg^-1^ nutrient *i*); *r_i_* = minimum nutrient *i* uptake to produce any grain (kg nutrient *i* ha^-1^).

The physiological efficiency (*PhE*) was calculated as follows (Sattari *et al.,* 2014):

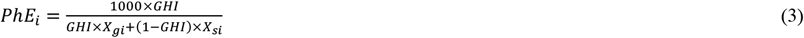

Where *GHI* = grain harvest index, *X_gi_* = mass fraction (g kg^-1^) of the nutrient *i* in the grain, *X_si_* = mass fraction (g kg^-1^) of the nutrient *i* in the stover. The GHI <0.40 values were considered as anomalies in the dataset as the crop might have suffered biotic and abiotic stresses other than nutrients (Hay, 1995); to guarantee accuracy they were excluded from this analysis.

The minimum uptake of the *i*^th^ nutrient to produce any grain (*r_i_*) was obtained from the minimum uptake of the *i*^th^ nutrient in the above ground biomass mass (kg ha^-1^) in the control plots after discarding all control plots with zero grain yield.

*Step 3 (relation between actual uptake and yield ranges)*: The principles used in QUEFTS at this stage are that the yield ranges are calculated between yield 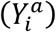 at maximum accumulation (*a*) and yield 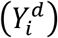 at maximum dilution (*d*), as functions of the actual uptake (*U_i_*) and the minimum uptake to produce any grain (*r_i_*):

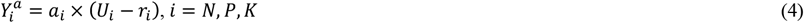

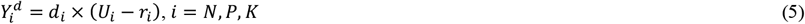

*Step 4 (combining yield ranges to ultimate yield estimates)*: in this final step yield ranges are combined for pairs of nutrients, and then the yields estimated for pairs of nutrients are averaged to obtain an ultimate yield estimate. The following equation was used to calculate yield (*Y_ij_*) for the pair of nutrients *i* and *j* (Sattari *et al.,* 2014):

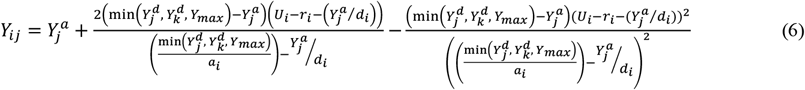

*i,j,k* = *N,P,K,* í ≠ *j* ≠ *k*; *Y_max_* = maximum potential yield (where 10,000 kg ha^-1^ was used in the study area).

The final and ultimate yield estimate (*Y_U_*) is calculated as the mean of the yield estimate of the pairs of nutrients:

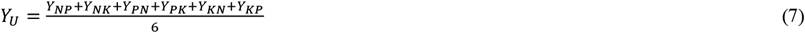

#### 2.4.2. Model validation

The root mean square error (RMSE), coefficient of determination (R^2^) and percent bias (PBIAS) were used to evaluate the precision and accuracy of QUEFTS model in predicting grain yield:

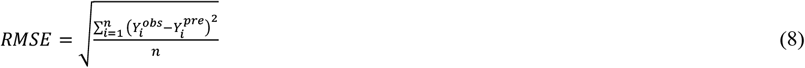

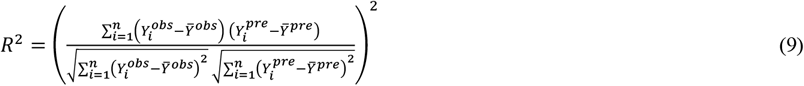

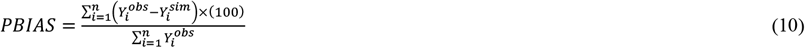

Where 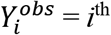 grain yield observed, *Ȳ^obs^* = mean of the observed grain yield, 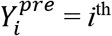 grain yield predicted by the QUEFTS model, *Ȳ^pre^* = mean of the predicted grain yield and *n* = number of observations.

The RMSE is an error index where the lower the value indicates better model performance (Moriasi *et al.,* 2007). The coefficient of determination (R^2^) estimates the combined dispersion against the single dispersion of the observed and predicted series (Krause and Boyle, 2005); it ranges between 0 and 1, where a value of 0 means no correlation at all and value of 1 means the dispersion of prediction is equal to that of observation. The optimal value of PBIAS is 0.00, with low-magnitude values indicating accurate model simulation. Positive values indicate model underestimation bias, and negative values indicate model overestimation bias (Gupta *et al.*, 1999).

## 3. Results

### 3.1 Soil characteristics of the experimental fields

There was a strong variability in most soil characteristics among the experimental fields across the two agro-ecological zones (NGS and SS) as indicated by wide range and high coefficient of variability (CV) values (Table 1). However, most of the studied parameters were significantly different between the two agro-ecological zones. Total organic carbon (OC_tot_), total nitrogen (N_tot_), Mg, Cu and available sulphur (S_av_) were larger in the NGS than in the SS. In contrast, pH, available phosphorus (P_av_), Mn and Fe were larger in the SS than in the NGS. In both agro-ecological zones, soils have a large sand content and are classified as loam in the NGS and sandy loam in the SS. The average soil pH is classified as moderately acidic (5.6-6.0) in the NGS and slightly acidic (6.1-6.5) in the SS. The average contents of OC_tot_ (<10 g kg^−1^), N_tot_ (< 0.10 g kg^−1^), B (< 0.79 mg kg^−1^) and ECEC (< 6.0 cmolc kg^−1^) in both agro-ecological zones fell within a low soil fertility condition according to the ratings of the Nigerian “National Special Programme on Food Security” NSPFS (2005) and of the ESU (1991) fertility classification of Nigerian Savanna soils. However, soil average P_av_ (7-20 mg kg^−1^), K (0.150.30 cmolc kg^−1^), Ca (2-5 cmolc kg^−1^), Mg (0.3-1.0 cmolc kg^−1^), Cu (0.21-2.0 mg kg^−1^) and S_av_ (5.1-20.0 mg kg^−1^) were of ‘moderate’ soil fertility status in both agro-ecological zones. High levels of Zn (> 2.0 mg kg^−1^), Mn (> 5.0 mg kg^−1^) and Fe (> 5.0 mg kg^−1^) were observed in the two agro-ecological zones.

**Table 1:** Selected physico-chemical properties of topsoil (0-20cm) of the experimental fields between the study agro-ecological zones

| Soil Properties | NGS |  | SS |  | P-Value |
| --- | --- | --- | --- | --- | --- |
|  | Mean (Range) | CV (%) | Mean (Range) | CV (%) |  |
| pH<sub>H2O</sub> (1:1) | 5.8 (4.8-7.2) | 8 | 6.2 (5.2-7.2) | 9 | <0.001** |
| OC<sub>tot</sub> (g kg<sup>-1</sup>) | 7.25 (2.44-15.45) | 36 | 5.01 (2.04-10.12) | 36 | <0.001** |
| N<sub>tot</sub> (g kg<sup>-1</sup>) | 0.47 (0.25-0.98) | 30 | 0.36 (0.17-0.66) | 36 | <0.001** |
| P<sub>av</sub> (mg kg<sup>-1</sup>) | 8.43 (0.64-31.77) | 82 | 16.54 (1.44-50.00) | 71 | <0.001** |
| Ca (cmol<sub>c</sub> kg<sup>-1</sup>) | 2.31 (0.28-9.78) | 43 | 2.54 (0.38-5.32) | 40 | 0.238 |
| Mg (cmol<sub>c</sub> kg<sup>-1</sup>) | 0.73 (0.07-1.99) | 41 | 0.59 (0.26-1.35) | 38 | 0.011* |
| K (cmol<sub>c</sub> kg<sup>-1</sup>) | 0.22 (0.06-1.35) | 78 | 0.24 (0.07-0.50) | 43 | 0.499 |
| Na (cmol<sub>c</sub> kg<sup>-1</sup>) | 0.08 (0.04-0.10) | 19 | 0.08(0.04-0.14) | 31 | 0.796 |
| EA (cmol<sub>c</sub> kg<sup>-1</sup>) | 0.04 (0.00-1.00) | 7 | 0.02 (0.00-0.15) | 8 | 0.307 |
| ECEC (cmol<sub>c</sub> kg<sup>-1</sup>) | 3.37 (1.23-11.06) | 34 | 3.45 (1.01-7.17) | 36 | 0.706 |
| Zn (mg kg<sup>-1</sup>) | 8.66 (0.83-69.06) | 87 | 9.43 (1.73-37.88) | 80 | 0.589 |
| Cu (mg kg<sup>-1</sup>) | 2.00 (0.76-5.12) | 47 | 1.52 (0.76-2.55) | 35 | 0.004** |
| Mn (mg kg<sup>-1</sup>) | 30.9 (3.71-158.46) | 65 | 45.76 (7.49-87.50) | 53 | <0.001** |
| Fe (mg kg<sup>-1</sup>) | 142.78 (43.36-327.18) | 57 | 207.22(122.87-439.14) | 34 | <0.001** |
| B (mg kg<sup>-1</sup>) | 0.03 (0.004-0.120) | 72 | 0.02 (0.003-0.100) | 112 | 0.528 |
| S<sub>av</sub> (mg kg<sup>-1</sup>) | 7.29 (4.55-11.70) | 20 | 6.25 (4.09-9.95) | 26 | <0.001** |
| Sand (%) | 45 (23-70) | 20 | 65 (47-77) | 11 | <0.001** |
| Silt (%) | 32 (13-59) | 22 | 19 (9-33) | 31 | <0.001** |
| Clay (%) | 23 (13-42) | 24 | 16 (12-23) | 18 | <0.001** |

NGS: Northern Guinea Savanna; SS: Sudan Savanna; OC<sub>tot</sub>: total organic carbon; N<sub>tot</sub>: total nitrogen; P<sub>av</sub>: available P; S<sub>av</sub>: available sulphur; EA: exchange acidity; ECEC: effective cation exchange capacity; CV: coefficient of variability.

### 3.2 Characteristics of grain yield and nutrient uptake of the experimental data

Nutrient application (NA) significantly affected all measured grain yield and nutrient uptake characteristics (Table 2). Maize grain yield, total dry matter, N and P uptake were consistently larger in the NPK+, NPK and -K nutrient application treatments than in the -P, -N and control, across the two agro-ecological zones (Fig. 3). Similar trend was observed for grain harvest index (GHI), K uptake and nutrient harvest indices (NHI, PHI, and KHI) except in the SS where the values of these variables for -P treatment were comparable with the values for NPK+, NPK and -K, respectively (Fig. 3). With an exception of plant P uptake (kg ha^−1^) and P harvest index (PHI), all the studied parameters for grain yield and nutrient uptake were significantly different between the agro-ecological zones (AEZ) (Table 2). Grain yield and total dry matter were on average largest in NGS (3.8 and 8.6 t ha^−1^) and smallest in SS (3.0 and 6.5 t ha^−1^) (Fig.3). Nitrogen (N) and K uptake were equally larger in the NGS (69.2 and 77.7 kg ha^−1^) than in the SS (52.1 and 60.1 kg ha^−1^) (Fig.3). In contrast, GHI, nitrogen harvest index (NHI) and potassium harvest index (KHI) were larger in the SS than in the NGS. There were few differences between the two variety groups (OPV and hybrid) (Table 2), with only GHI, NHI and PHI being larger in the OPV than in the hybrid variety group (Fig. 4). However, significant interaction among variety group and agro-ecological zone on GHI and N, P and K harvest indices were also observed (Table 2). The GHI was comparable between the two variety groups in the NGS, while in the SS an OPV had larger GHI (0.49) than the hybrid variety (0.41) (Fig. 4). Largest N, P and K harvest indices (NHI, PHI and KHI) were observed in OPV in the SS zone. Because of a few statistical differences between variables of the two variety groups, the datasets from the two groups were combined in the parameterization of the QUEFTS model.

**Fig. 3:**
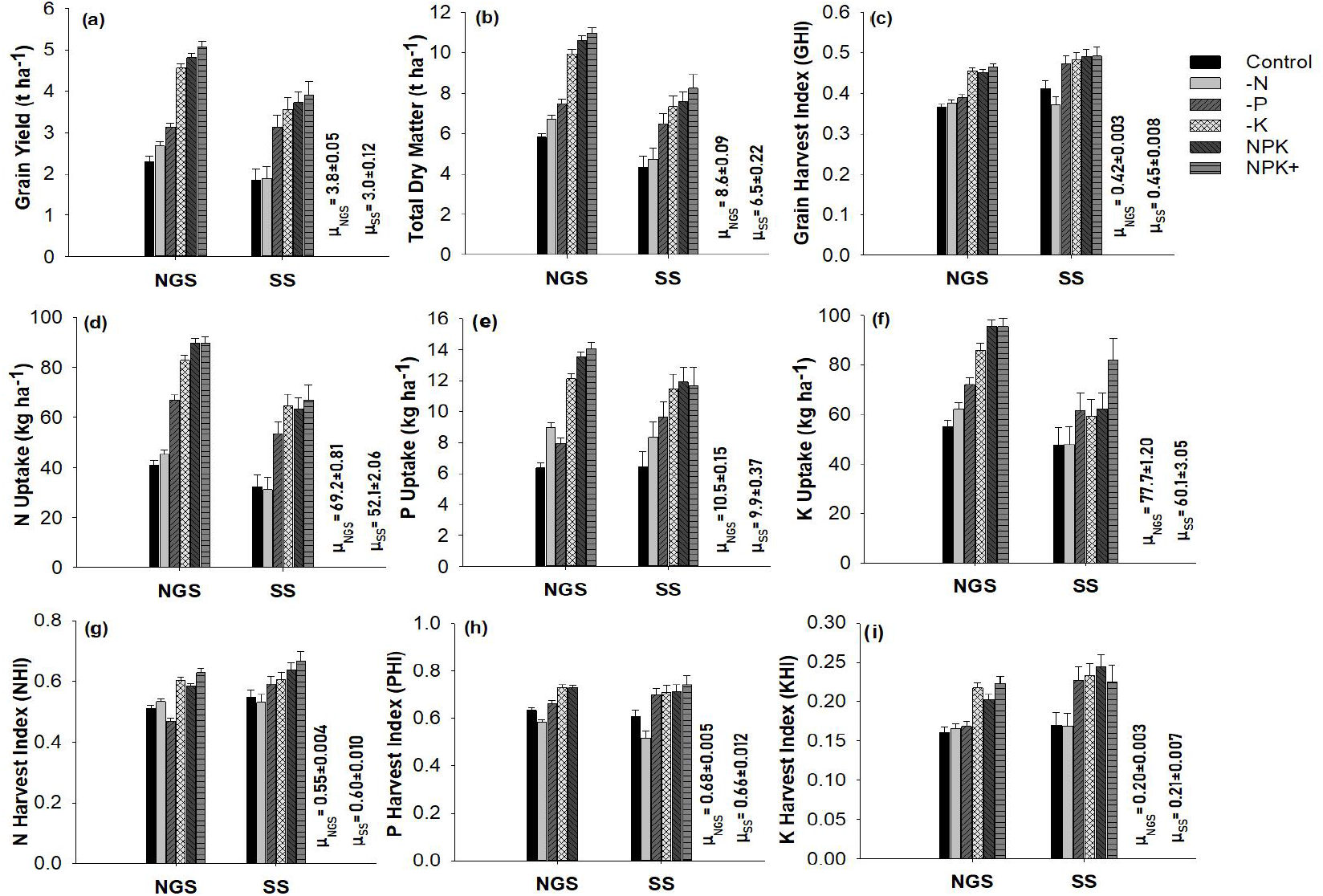
Effects of nutrient application (NA) across agro-ecological zones (AEZ) on (**a**) maize grain yield (**b**) total dry matter (**c**) grain harvest index (d) N uptake (**e**) P uptake (**f**) K uptake (**g**) N harvest index (**h**) P harvest index and (**i**) K harvest index. Error bars are standard error of means; NGS: Northern Guinea Savanna; SS: Sudan Savanna; μNGS: mean NGS, μSS = mean SS.

**Fig. 4:**
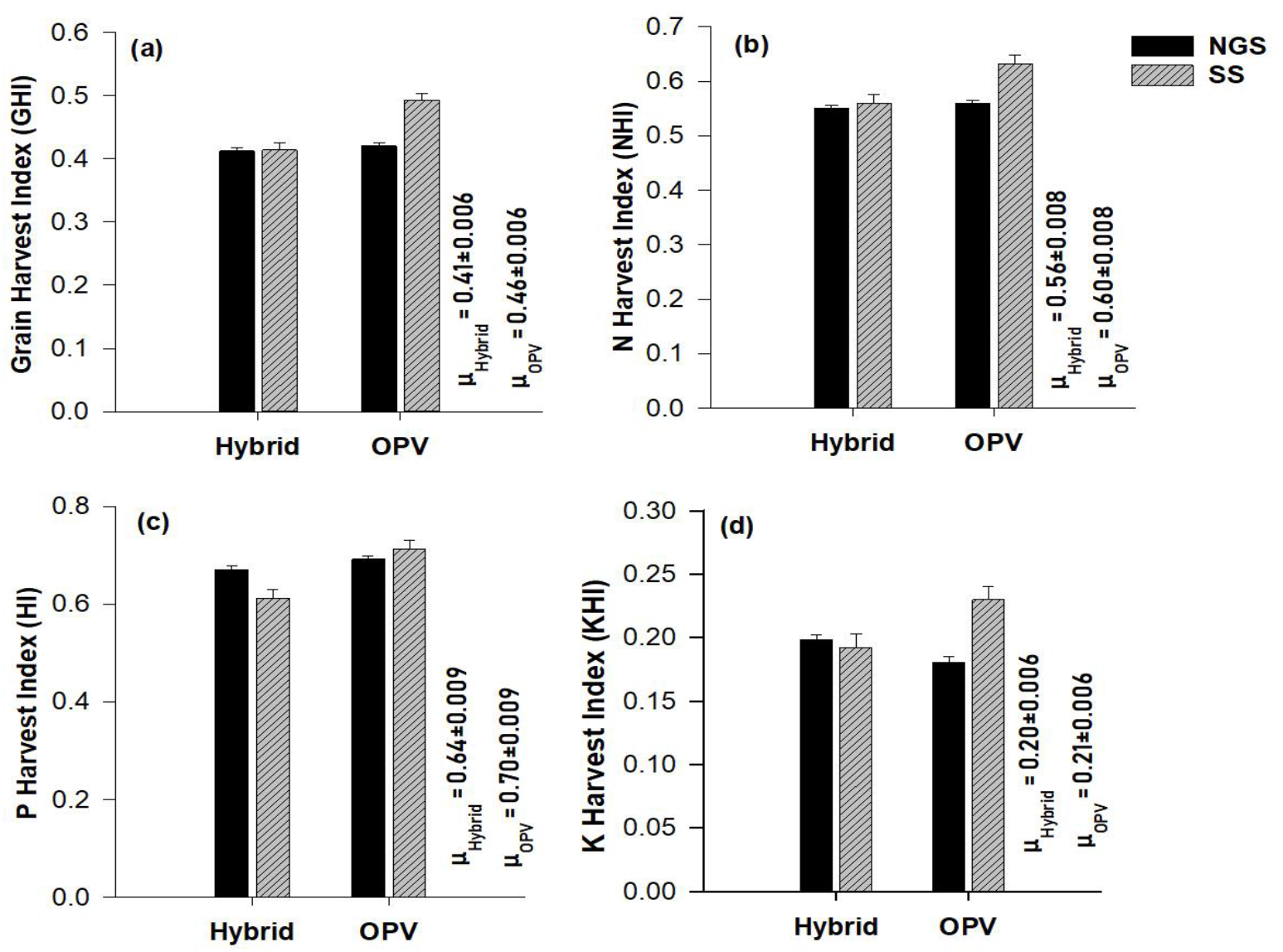
Effects of variety group (VG) across agro-ecological zones (AZE) on (**a**) grain harvest index (**b**) N harvest index (**c**) P harvest index (**d**) K harvest index. Error bars are standard error of means; NGS: Northern Guinea Savanna; SS: Sudan Savanna; OPV: open pollinated variety; hybrid: hybrid variety; μ_Hybrid_: mean hybrid, μ_OPV_ = mean OPV.

**Table 2:** Probability of F values (P-Value) of response of grain yield and nutrient uptake parameters to nutrient application (NA), agro-ecological zones (AEZ) and variety group (VG) of the experimental data.

| Parameter | Main Effect |  |  | Interaction Effect |  |  |  |
| --- | --- | --- | --- | --- | --- | --- | --- |
|  | NA | AEZ | VG | NA x AEZ | NA x VG | AEZ x VG | NA x AEZ x VG |
| Grain yield (t ha<sup>-1</sup>) | <0.001*** | <0.001*** | 0.634 | 0.074 | 0.543 | 0.571 | 0.893 |
| Total dry matter (t ha<sup>-1</sup>) | <0.001*** | <0.001*** | 0.103 | 0.074 | 0.734 | 0.176 | 0.820 |
| Grain Harvest index | <0.001*** | <0.001*** | <0.001*** | 0.070 | 0.174 | <0.001*** | 0.429 |
| Plant N uptake (kg ha<sup>-1</sup>) | <0.001*** | <0.001*** | 0.997 | 0.173 | 0.935 | 0.667 | 0.882 |
| Plant P uptake (kg ha<sup>-1</sup>) | <0.001*** | 0.119 | 0.632 | 0.077 | 0.678 | 0.579 | 0.894 |
| Plant K uptake (kg ha<sup>-1</sup>) | <0.001*** | <0.001*** | 0.293 | 0.105 | 0.679 | 0.065 | 0.477 |
| N harvest index | <0.001*** | <0.001*** | <0.001*** | 0.016* | 0.229 | <0.001*** | 0.168 |
| P harvest index | <0.001*** | 0.129 | <0.001*** | 0.311 | 0.125 | <0.001*** | 0.373 |
| K harvest index | <0.001*** | <0.001*** | 0.194 | 0.183 | 0.096 | <0.001*** | 0.421 |

### 3.3 QUEFTS model parameterization

#### 3.3.1 Indigenous soil nutrient supply and fertilizer recovery efficiency

The relations between indigenous soil N, P, and K supply (calculated as the uptake of the given nutrient in the respective omission plots) and soil properties were not well described with the QUEFTS’ default equations (Table 3) both in the NGS and the SS as could be derived from the relatively small R^2^ values. Consequently, new and better fitting equations of indigenous soil N, P and K supply were derived for the NGS and the SS (Table 3). In addition to OC_tot_, an explanatory soil property for indigenous N supply in the default QUEFTS model, positive contributions from N_tot_ and Mg led to new explanatory soil properties in the NGS. In the SS, N_tot_, Cu, and Mn contributed positively as the explanatory soil properties for the indigenous supply of N. Unlike the default QUEFTS model, OC_tot_ did not have significant influence on the indigenous supply of P and K in both agro-ecological zones. Indigenous soil supply of P was explained with only pH and P_av_ in the NGS, and with pH, P_av_ and Cu in the SS, in all cases with a positive contribution. Exchangeable potassium (K), ECEC and clay content were the significant explanatory soil properties for K indigenous soil supply in the NGS, with K and ECEC having a positive effect, and clay content having a negative effect. In the SS, OC_tot_, K, Mn, pH and silt contents were explaining indigenous K-supply from soil, in a positive way for the first three and negatively for the latter two (Table 3).

**Table 3:** Default and improved/parameterized indigenous maize N, P and K supply equations for the Northern Guinea Savanna (NGS) and the Sudan Savanna (SS)

| Nutrient | Calibrated | QUEFTS Default (Janssen et al., 1990) |
| --- | --- | --- |
| NGS |  |  |
| N | $S_N = -28.87 + 0.40 \text{ OC}_{\text{tot}} (\text{g kg}^{-1}) + 119.90 \text{ N}_{\text{tot}} (\text{g kg}^{-1}) + 0.31 \text{ Silt } (\%) + 7.56 \text{ Mg } (\text{cmol}_c \text{ kg}^{-1})$<br>(**R<sup>2</sup>=0.60**) | $S_N = 22.80 + 2.54 \text{ OC}_{\text{tot}} (\text{g kg}^{-1})$<br>(**R<sup>2</sup>=0.11**) |
| P | $S_P = -12.16 + 2.71 \text{ pH} + 0.71 \text{ P}_{\text{av}} (\text{mg kg}^{-1})$<br>(**R<sup>2</sup>=0.61**) | $S_P = 5.46 - 0.22 \text{ OC}_{\text{tot}} (\text{g kg}^{-1}) + 0.72 \text{ P}_{\text{av}} (\text{mg kg}^{-1})$<br>(**R<sup>2</sup>=0.57**) |
| K | $S_K = 25.56 - 1.67 \text{ Clay } (\%) + 231.21 \text{ K } (\text{cmol}_c \text{ kg}^{-1}) + 13.54 \text{ ECEC } (\text{cmol}_c \text{ kg}^{-1})$<br>(**R<sup>2</sup>=0.62**) | $S_K = 37.53 - 1.60 \text{ OC}_{\text{tot}} (\text{g kg}^{-1}) + 248.05 \text{ K } (\text{cmol}_c \text{ kg}^{-1})$<br>(**R<sup>2</sup>=0.46**) |
| SS |  |  |
| N | $S_N = -28.79 + 90.11 \text{ N}_{\text{tot}} (\text{g kg}^{-1}) + 5.53 \text{ Cu } (\text{mg kg}^{-1}) + 0.22 \text{ Mn } (\text{mg kg}^{-1})$<br>(**R<sup>2</sup>=0.62**) | $S_N = 24.87 + 0.61 \text{ OC}_{\text{tot}} (\text{g kg}^{-1})$<br>(**R<sup>2</sup>=0.03**) |
| P | $S_P = -8.59 + 0.98 \text{ pH} + 0.40 \text{ P}_{\text{av}} (\text{mg kg}^{-1}) + 2.85 \text{ Cu } (\text{mg kg}^{-1})$<br>(**R<sup>2</sup>=0.69**) | $S_P = 3.29 - 0.11 \text{ OC}_{\text{tot}} (\text{g kg}^{-1}) + 0.31 \text{ P}_{\text{av}} (\text{mg kg}^{-1})$<br>(**R<sup>2</sup>=0.56**) |
| K | $S_K = 272.15 - 49.75 \text{ pH} + 7.30 \text{ OC}_{\text{tot}} (\text{g kg}^{-1}) - 1.27 \text{ Silt } (\%) + 324.89 \text{ K } (\text{cmol}_c \text{ kg}^{-1}) + 0.48 \text{ Mn } (\text{mg kg}^{-1})$<br>(**R<sup>2</sup>=0.76**) | $S_K = 39.13 - 2.50 \text{ OC}_{\text{tot}} (\text{g kg}^{-1}) + 237.21 \text{ K } (\text{cmol}_c \text{ kg}^{-1})$<br>(**R<sup>2</sup>=0.36**) |

OC<sub>tot</sub>: total organic carbon; N<sub>tot</sub>: total nitrogen; P<sub>av</sub>: available P; ECEC: effective cation exchange capacity; S<sub>N</sub>, S<sub>P</sub> and S<sub>K</sub> are soil indigenous supplies in kg ha<sup>-1</sup> of maize crop-available N, P, and K, respectively.

Both the newly parameterized and default QUEFTS average fertilizer recovery efficiencies are shown in Table 4. The fertilizer recovery fractions of N, P and K were generally larger in the NGS than in the SS (Table 4). In both agro-ecological zones recovery efficiencies of N were smaller than the QUEFTS default value of 0.50. The average P and K recovery efficiencies were larger than the QUEFTS default efficiency values of 0.10 and 0.50, respectively in the NGS. On the contrary, the average P and K recovery efficiencies were smaller than the QUEFTS default values in the SS (Table 4).

**Table 4:** Default and newly parameterized values of average fertilizer recovery efficiency (*R_i_*); physiological efficiency at maximum accumulation of nutrient (*a_i_*) and maximum dilution of nutrient (*d_i_*); and minimum uptake required (*r_i_*) to produce any grain of N, P and K in the above-ground dry matter of maize in the Northern Guinea (NGS) and the Sudan Savanna (SS)

| Coefficients | Nutrients | Default QUEFTS Model<br>(Janssen et al.,1990) | NGS | SS |
| --- | --- | --- | --- | --- |
| Average fertilizer recovery fraction<br>“$R_i$” | N | 0.50 | 0.42 | 0.32 |
|  | P | 0.10 | 0.16 | 0.08 |
|  | K | 0.50 | 0.54 | 0.37 |
| Physiological efficiency at<br>maximum accumulation of the<br>nutrient “$a_i$” (kg grain kg<sup>-1</sup> nutrient) | N | 30 | 35 | 32 |
|  | P | 200 | 200 | 164 |
|  | K | 30 | 25 | 24 |
| Physiological efficiency at<br>maximum dilution of the nutrient<br>“$d_i$” (kg grain kg<sup>-1</sup> nutrient) | N | 70 | 79 | 79 |
|  | P | 600 | 527 | 528 |
|  | K | 120 | 117 | 136 |
| Minimum nutrient uptake to<br>produce any grain “$r_i$” (kg ha<sup>-1</sup>) | N | 5.0 | 4.0 | 6.1 |
|  | P | 0.4 | 0.5 | 0.8 |
|  | K | 2.0 | 4.5 | 7.3 |

#### 3.3.2 Physiological nutrient efficiency and minimum nutrient uptake to produce any grain

The relations between grain yield and nutrient uptake showing boundary lines of physiological efficiency (*PhE*) of nutrients at maximum accumulation (*a*) and maximum dilution (*d*) are presented in Fig. 5. Across the two agro-ecological zones, the coefficients *a* for N, P and K were overall close to the QUEFTS default values (Table 4). The only exception was in the SS where coefficient *a* for P was lower than the QUEFTS standard value. The *d* coefficients for N between the NGS and SS were comparable, but larger than the QUEFTS default value. In contrast, the *d* coefficients for P between the two agro-ecological zones were comparable, but lower than the QUEFTS default value. The *d* coefficient for K in the NGS was close to the QUEFTS default value, but these values were lower than the value observed in the SS. The values for the minimum nutrient uptake coefficient (*r*) of N, P and K were 4.0, 0.5 and 4.5 kg ha^−1^ for the NGS; and 6.1, 0.8 and 7.3 kg ha^−1^ for the SS, respectively (Table 4). Across the two agro-ecological zones, the *r* coefficient values for all the three nutrients (N, P, and K) were larger than the QUEFTS default values, except *r* coefficient for the N in the NGS, which was slightly smaller than the QUEFTS default coefficient. However, the *r* coefficient values of the three nutrients were smaller in the NGS than in the SS.

**Fig. 5:**
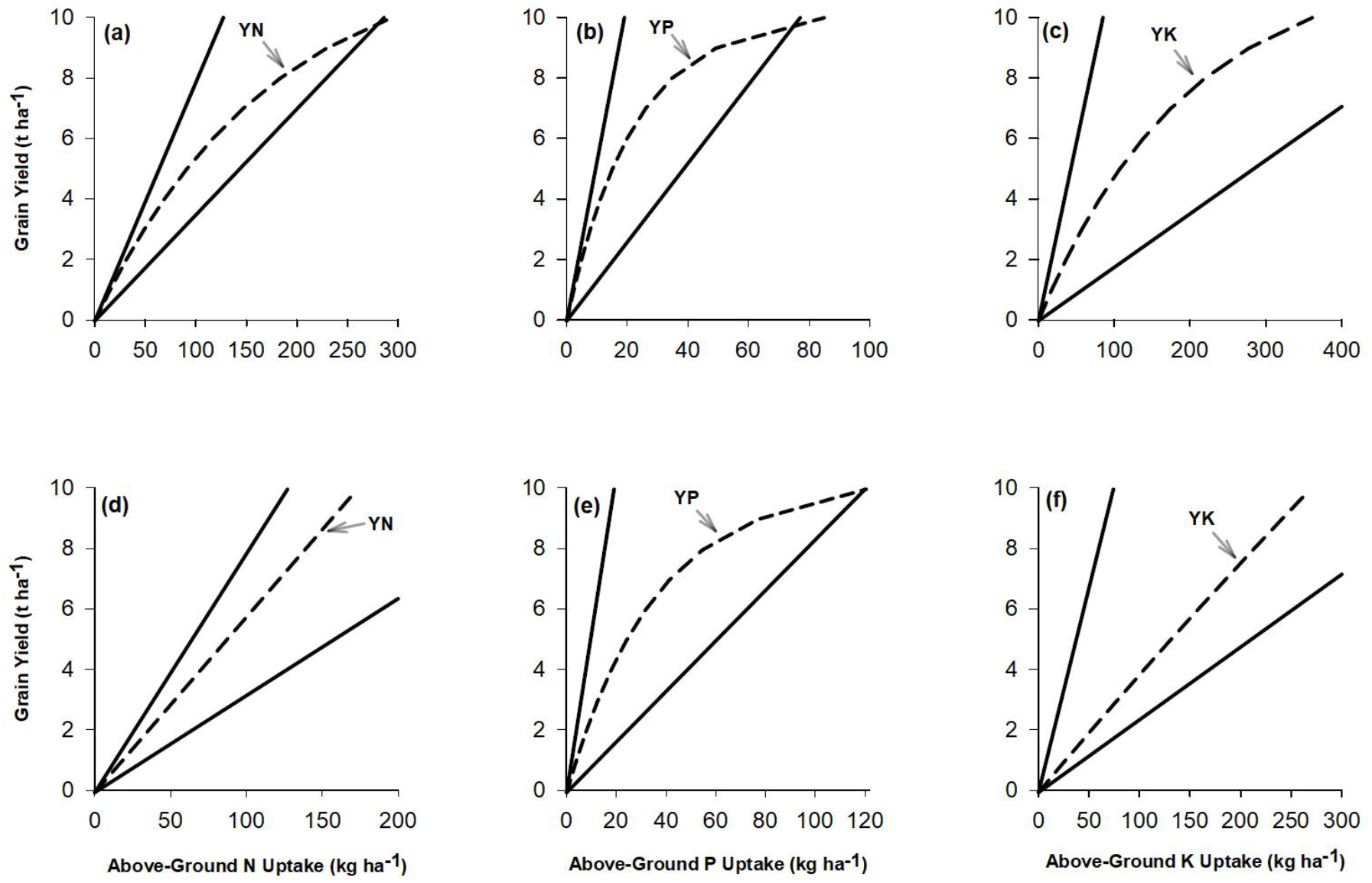
The balanced maize N, P and K uptake requirements (YN, YP and YK) for maximum yield potentials set at 10 t ha^−1^ simulated by the parameterized-QUEFTS model for Northern Guinea Savanna (a-c) and Sudan Savanna (d-f). The upper and lower lines indicate yields with maximum dilution and maximum nutrient accumulation, respectively.

### 3.4 Balanced nutrient uptake requirements

The QUEFTS model predicts a linear relationship between grain yield and above-ground nutrient uptake until yield reaches about 50-60% of the yield potential fixed at 10 t ha^−1^ for the NGS and the SS, respectively (Fig. 5). As the target yield gets closer to the potential yield, *PhE* decreases significantly (Table 5). The parametrized QUEFTS model estimated a balanced uptake of 19.4 kg N, 3.3 kg P and 23.0 kg K in the above-ground parts per tonne of maize grain yield when the grain yield reached 60% (6 t ha^−1^) of the maize potential yield in the NGS (Table 5). The corresponding *PhE* was 51.6 kg grain kg^−1^ N, 302.9 kg grain kg^−1^ P and 43.5 kg grain kg^−1^ K. Likewise, in the SS an uptake of 17.3 kg N, 5.3 kg P and 26.2 kg K was required per tonne of grain yield at 60% of the potential yield (Table 5); the corresponding *PhE* was 57.9 kg grain kg^−1^ N, 190.6 kg grain kg^−1^ P and 38.2 kg grain kg^−1^ K. It follows that the optimal N, P & K ratios in the above-ground dry matter at 60% of the maize potential yield are 5.9:1:7.0 for the NGS and 3.3:1:4.9 for the SS. These results show that the QUEFTS model predicts larger P and K uptake requirements for a balanced nutrition at 60% of the potential yield in the SS than in the NGS, while an opposite trend was observed for N requirements between the agro-ecological zones.

**Table 5:**
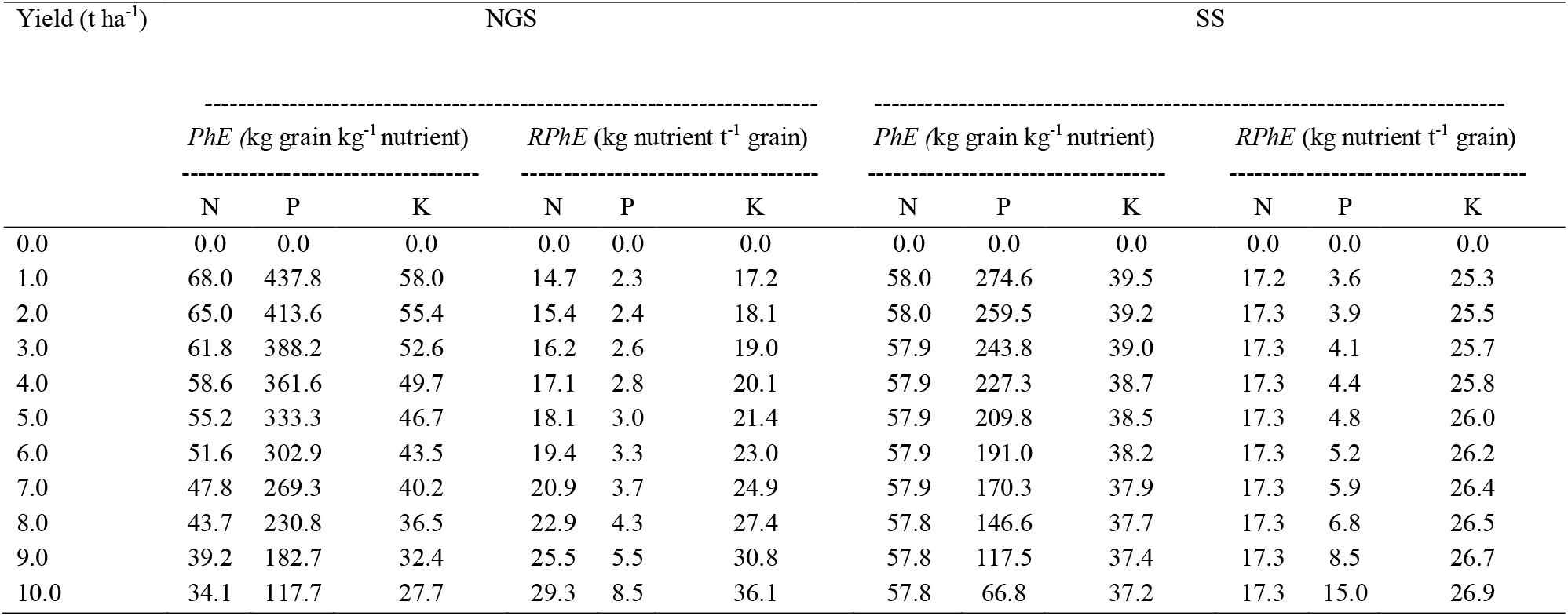
Maize physiological efficiency (*PhE*) and reciprocal physiological efficiency (*RPhE*) of N, P, and K simulated by the QUFETS model to achieve yield targets with maximum yield potential set at 10 t ha^−1^ for the Northern Guinea Savanna (NGS) and the Sudan Savanna (SS)

| Yield (t ha<sup>-1</sup>) | NGS |  |  |  |  |  | SS |  |  |  |  |  |
| --- | --- | --- | --- | --- | --- | --- | --- | --- | --- | --- | --- | --- |
|  | *PhE* (kg grain kg<sup>-1</sup> nutrient) |  |  | *RPhE* (kg nutrient t<sup>-1</sup> grain) |  |  | *PhE* (kg grain kg<sup>-1</sup> nutrient) |  |  | *RPhE* (kg nutrient t<sup>-1</sup> grain) |  |  |
|  | N | P | K | N | P | K | N | P | K | N | P | K |
| 0.0 | 0.0 | 0.0 | 0.0 | 0.0 | 0.0 | 0.0 | 0.0 | 0.0 | 0.0 | 0.0 | 0.0 | 0.0 |
| 1.0 | 68.0 | 437.8 | 58.0 | 14.7 | 2.3 | 17.2 | 58.0 | 274.6 | 39.5 | 17.2 | 3.6 | 25.3 |
| 2.0 | 65.0 | 413.6 | 55.4 | 15.4 | 2.4 | 18.1 | 58.0 | 259.5 | 39.2 | 17.3 | 3.9 | 25.5 |
| 3.0 | 61.8 | 388.2 | 52.6 | 16.2 | 2.6 | 19.0 | 57.9 | 243.8 | 39.0 | 17.3 | 4.1 | 25.7 |
| 4.0 | 58.6 | 361.6 | 49.7 | 17.1 | 2.8 | 20.1 | 57.9 | 227.3 | 38.7 | 17.3 | 4.4 | 25.8 |
| 5.0 | 55.2 | 333.3 | 46.7 | 18.1 | 3.0 | 21.4 | 57.9 | 209.8 | 38.5 | 17.3 | 4.8 | 26.0 |
| 6.0 | 51.6 | 302.9 | 43.5 | 19.4 | 3.3 | 23.0 | 57.9 | 191.0 | 38.2 | 17.3 | 5.2 | 26.2 |
| 7.0 | 47.8 | 269.3 | 40.2 | 20.9 | 3.7 | 24.9 | 57.9 | 170.3 | 37.9 | 17.3 | 5.9 | 26.4 |
| 8.0 | 43.7 | 230.8 | 36.5 | 22.9 | 4.3 | 27.4 | 57.8 | 146.6 | 37.7 | 17.3 | 6.8 | 26.5 |
| 9.0 | 39.2 | 182.7 | 32.4 | 25.5 | 5.5 | 30.8 | 57.8 | 117.5 | 37.4 | 17.3 | 8.5 | 26.7 |
| 10.0 | 34.1 | 117.7 | 27.7 | 29.3 | 8.5 | 36.1 | 57.8 | 66.8 | 37.2 | 17.3 | 15.0 | 26.9 |

### 3.5 QUEFTS model validation

Fig. 6 shows the comparison between observed and parameterized QUEFTS predicted maize grain yields for the NGS and the SS. There was a good agreement between grain yields predicted by the parameterized QUEFTS model and those observed from the field experiment in both the NGS and the SS based on reasonably high R^2^ (0.70 and 0.86, respectively) and small RMSE values (1.01 and 0.79, respectively) (Fig. 7). However, the model showed a small overestimation bias in the NGS (PBIAS = −10.0%) and a small underestimation bias in the SS (PBIAS = 13.0%).

**Fig. 6:**
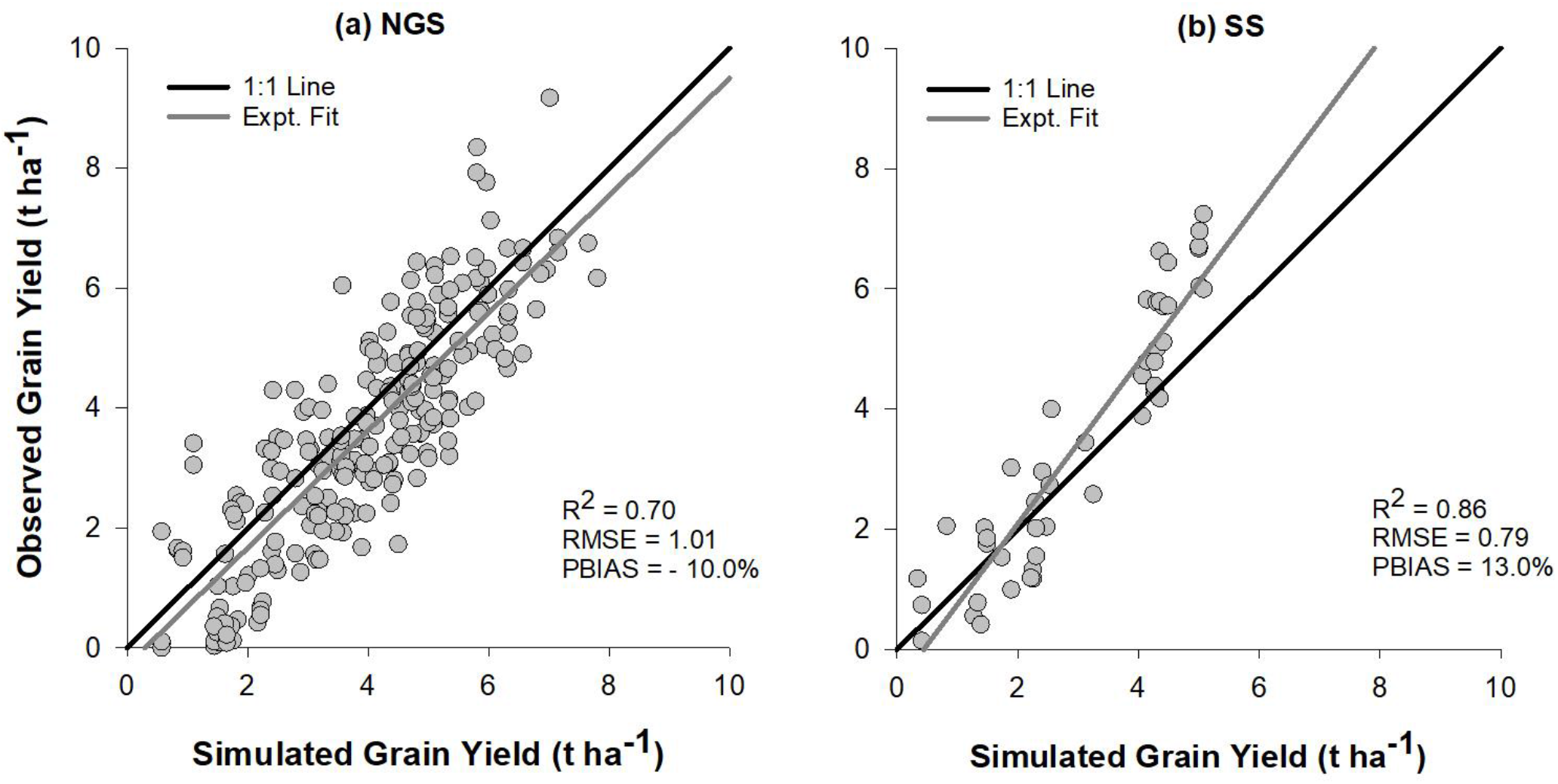
Relation between the observed and parameterized QUEFTS simulated maize grain yield for (a) Northern Guinea Savanna “NGS” (b) Sudan Savanna “SS”.

## 4. Discussion

### 4.1 Soil characteristics of the experimental fields

The larger OC_tot_, N_tot_ and S_av_ in NGS is associated with larger soil clay content as well as longer rainy season which resulted in greater vegetative biomass and litterfall than in SS. In contrast, relatively less rainfall in the SS resulted in higher average pH values compared to the NGS. High rainfall increases the potential for leaching of cations (especially Ca and Mg) and poor soil aeration, which often decreases soil pH. The higher OC_tot_, N_tot_ and S_av_, and lower pH in the NGS than in the SS has also been observed by Daudu et al. (2018). Northern Nigerian Savanna soils are developed from aeolian materials and pre-Cambrian basement complex rocks (such as granite, schist and sandstone) (Bennett, 1980) and this resulted in a large sand fraction in the surface soils of both the NGS and the SS. Moreover, Malgwi et al. (2000) and Voncir et al. (2008) have reported that sorting of soil material due to clay eluviation and wind erosion as additional factors leading to large sand content of the surface soils of the Northern Nigerian Savanna. The overall low levels of OC_tot_, N_tot_, B and ECEC in both agro-ecological zones have been related to two principal factors: (i) the type of parent material and intensive weathering of the soils with small mineral reserve necessary for inherent nutrient recharge; and (ii) intensive cultivation of the soils with inadequate (unbalanced and insufficient external inputs) nutrient management including burning or complete removal of crop residues (Jones and Wild, 1975; Manu et al., 1991; Smaling et al., 1991; Kwari et al., 2011). Although past studies like Ekeleme et al. (2014), Kamara et al. (2014) and Shehu et al. (2015) have reported low average P_av_ in some parts of the study area, the moderate average P_av_ content observed in the two agro-ecological zones in this study can be explained by residual effects of previous P applications. Some fractions of P applied through fertilizer not taken up by the current crop due to temporary fixation in the soil can be released gradually to the succeeding crop (Janssen et al., 1987). The moderate to high levels of exchangeable cations (Ca, Mg and K) are not surprising as most soils are developed from basement complex rocks which contains large content of these cations. Møberg and Esu (1991) also reported an appreciable presence of K-bearing feldspar minerals in sand and silt particles of Savanna soils of Nigeria. Similarly, deficiency of the studied micronutrients (Fe, Mn, Cu and Zn) is unlikely to occur in the study fields due to relatively acidic reaction of the soils. Only at pH above 7.5 does the availability of these micronutrients becomes significantly limited owing to the formation of oxides, hydroxides and carbonates (Sillanpää, 1982).

### 4.2 Characteristics of grain yield and nutrient uptake of the experimental data

The minimal response of grain yield, total dry matter, GHI and N, P and K uptake in control, -N and -P relative to the NPK+, NPK and -K treatments across the two agro-ecological zones indicates N and P as the major nutrients limiting growth and yield response of maize. Nitrogen deficiency has been recognized as the most limiting factor for cereal production in vast areas of SSA including in the Nigerian Savanna (Vanlauwe et al., 2011). Soil N can be depleted rapidly by maize, especially when yields are high and stover is exported (Kamara, 2017). The widespread N deficiency in the study area can be attributed to small soil organic matter contents (indicated by small OC_tot_) resulting from inherent poor soil fertility and continuous cropping with inadequate and imbalanced N fertilizer or manure applications. Adediran and Banjoko (1995) reported P as among the most maize yield limiting nutrients in the Nigerian Savanna. Nigerian soils, particularly the highly weathered ones, have small indigenous P contents and often a large P sorption capacity (Osemwotai et al., 2005). Combined application of balanced fertilizers with manure and rotation of cereal crops with legumes through integrated soil fertility management principles (ISFM) (Vanlauwe et al., 2010) can assist farmers in the study area to improve soil N and P status. The lack of a significant increase in grain yield due the addition of secondary macronutrients (S, Ca and Mg) and micronutrients (Zn and B) suggest that these nutrients are not significantly limiting maize yield in the studied area. A significant extra yield increases due to the addition of the secondary macronutrients and micronutrients was observed in only 7 fields (Shehu et al., 2018). The larger grain yield and total dry matter in the NGS compared with the SS could be explained by the amount of rainfall, as the larger relative rainfall amount and duration in the NGS favoured more maize biomass production than in the SS.

#### 4.3.1 Indigenous soil nutrient supply and fertilizer recovery efficiency

The newly developed supply functions for indigenous soil N, P and K in both agroecological zones explained a minimum of 60% variation in soil characteristics among the studied fields. The unexplained variation can be attributed to the differences in rate of mineralization, in leaching losses and in soil moisture availability, etc. (Barber, 1995). These are complex factors to integrate into a simple empirical indigenous nutrient supply equation (Tabi et al., 2008). Going beyond the default QUEFTS model, total nitrogen (N_tot_) represents a more apt explanatory variable for the indigenous soil supply of nitrogen (S_N_) rather than the conventional OC_tot_, especially in the SS. Nitrogen mineralization in soil is indeed directly related to microbial activity and organic matter inputs, which are influenced by a combination of several physical, biological and chemical factors in the soil system (He, 2014). Hence, it is no surprise that OC_tot_ does not always provide the best proxy for N-availability in the soil. Samaké (2003) also reported OC_tot_ did not statistically influence indigenous supply of N, P and K to pearl millet in the similar soil conditions in Mali. Other nutrients like Mg in the NGS and Cu and Mn in the SS have shown to enhance the S_N_, emphasizing the importance of these (among other) nutrients in optimizing N supply. Magnesium together with N are involved in controlling processes of photosynthesis, assimilate production and partitioning among plant parts, which makes Mg a critical player in plant N uptake and utilization (Gastal and Lemaire, 2002; Shaul, 2002; Cakmak and Kirkby, 2008). Aulakh and Malhi (2005) reported that Mn uptake in plant can be stimulated by either or both forms of available N, the nitrate form may lead to a larger uptake of Mn and ammonium form may increase the bioavailability of Mn through acidification (Senbayram et al., 2015). The effect of pH on indigenous soil supply of P (S_P_) in both agro-ecologies corroborates the findings of Janssen et al. (1990). Most of the studied fields have acidic pH values, at this condition a unit decrease in pH level increases the potential of conversion of available phosphorus into a less soluble form through reacting with Al and Fe. Past studies indicated amount and type of clay as the dominant factor affecting the availability and supply of K in soil as observed for indigenous soil supply of potassium (S_K_) in the NGS. Fixation of K is larger in clayey than in sandy textured soils, and K fixation is largest in vermiculites and mica-clay minerals and smallest in montmorillonites and kaolinites (Sardi and Csitari, 1998; Conti et al., 2001).

Favourable combinations of adequate rainfall and low night temperatures makes the NGS more suitable for maize production than the SS (Badu-Apraku et al., 2015), this translates into the larger N, P and K fertilizer recovery efficiencies observed in the NGS. Despite in overall N and P recovery efficiency (R_N_ and R_P_) fell below the default QUEFTS values across the two agro-ecologies, but the values in the NGS are close to the result obtained by Saïdou et al. (2003) of 0.40 and 0.14 for N and P, respectively in the Southern Benin. In the same way, the recovery efficiency of K (R_K_) in the SS is in agreement with 0.40 reported in the Southern Benin by the same Saïdou et al. (2003). However, the R_P_ of both NGS and SS is smaller than the value of 0.24 observed by Tabi et al. (2008) in some part of the Northern Nigeria. This suggest that effective results which optimize fertilizer recovery efficiency figures can be obtained exclusively if site-specific nutrient recommendations using balanced nutrient requirements are complemented with the right source, time and place of fertilizer application, and subject to appropriate agronomic practices.

#### 4.3.2 Boundary line coefficients for physiological efficiency of nutrients and minimum nutrient uptake to produce any grain

The borderline coefficients *a* and *d* for physiological nutrient efficiency of this study across the two agro-ecological zones are larger than in the analysis of Saïdou et al. (2003) in the Southern Benin (20 and 40 kg grain kg^−1^ N, 110 and 270 kg grain kg^−1^ P, 25 and 90 kg grain kg^−1^ K) except *a* coefficient for K that are similar. Equally, Tabi et al. (2008) observed smaller *a* and *d* borderline physiological efficiency for N and P in some part of Northern Nigeria (21 and 71 kg grain kg^−1^ N, 97 and 600 kg grain kg^−1^ P) except *d* coefficient for P that is larger compared with the values of this study. Saïdou et al. (2003) and Tabi et al. (2008) have attributed the smaller physiological efficiencies in their studies to smaller grain harvest indices. Therefore, the larger values of physiological efficiencies in this study proved to be the result of large grain harvest indices. As explained earlier under sub-section 2.4.1, grain harvest indices less than 0.40 were considered as anomalies in the dataset as the crop might have suffered biotic and abiotic stresses other than nutrients (Hay, 1995); to guarantee precision were excluded as similarly performed by Liu et al. (2006), Xu et al. (2013), among others.

The significant difference between the minimum uptake requirement to produce any grain (*r*) observed in this study and the QUEFTS default values emphasizes the importance for recalibration of this parameter which has not been considered in most previous QUEFTS parameterization and calibration studies.

### 4.4 Balanced nutrient uptake requirements

Balanced nutrient plant uptake requirement can provide guidance for amount of fertilizer to be applied to achieve a desirable yield and for an efficient maintenance of soil fertility, as at least the nutrients removed or harvested in the above ground plant dry matter must be returned to the soil. The balanced nutrient uptake requirements predicted by QUEFTS in this study are comparable to values of 20.0 kg N, 4.5 kg P, 18.0 kg K reported for a tonne of maize grain in similar environmental and soil conditions in Zimbabwe (Piha, 1993). However, the higher balanced K uptake ratio in the above-ground matter relative to N as predicted by the parameterized QUEFTS in this study across the two agro-ecologies does not support the findings of most previous studies which have reported higher N uptake ratio compared to K. This trend was not surprising as most of the study fields have moderate to high K content in addition to the amount K fertilizer applied of 40-50 kg K ha^−1^. This led to luxury uptake of K especially in the maize stover evidenced by a small K harvest index (KHI). The moderate to high K content of the soils could be linked to an appreciable amount of K-bearing feldspar minerals in the sand and silt particles in the study area (Møberg and Esu, 1991) and the residual effect of previous K fertilizer applications. The supply of available K in soil is strongly dependent upon the type and amount of K-bearing minerals. In the K-feldspars, K is structurally bound in the crystal lattice (structural K) and is only released into the soil solution through weathering (Øgaard and Krogstad, 2005).

### 4.5 QUEFTS model validation

The close agreement between the parametrized QUEFTS simulated and observed yields shows that the parameterized QUEFTS model can be used to calculate balanced nutrient requirements and site- or area-specific fertilizer recommendations to optimize maize yield in the Northern Nigerian Savanna. The QUEFTS model however assumes that other biophysical factors apart from nutrients such as moisture, temperature, pests, diseases and management are non-limiting. As these factors are hard to optimize in on-farm field experiments, this may account for the under- and over-estimation bias obtained with the parameterized QUEFTS model in the SS and the NGS, respectively. To guarantee precision, the under- and overestimation percent bias in the SS and NGS, respectively should be considered and adjusted at the final and ultimate yield estimate (*Y_U_*) stage in the parameterized QUEFTS model.

## 5. Conclusion

The present study resulted in the parameterization and validation of the QUEFTS model to arrive at balanced nutrient requirements and site-specific fertilizer recommendations for maize in the Northern Nigerian Savanna. This was based on data from on-farm nutrient omission trials conducted across potential maize production sites covering two agro-ecological zones; i. e. the Northern Guinea Savanna (NGS) and the Sudan Savanna (SS). There were considerable differences in soil and nutrient uptake characteristics between the NGS and the SS. The relations between indigenous soil N, P, and K supply and soil properties were not well described with the QUEFTS default equations in both agro-ecological zones, consequently new and better fitting equations were derived. The coefficients *a* and *d* of N, P, and K for the QUEFTS model were 35 and 79, 200 and 527, and 25 and 117 kg grain kg^−1^ nutrient for the NGS and 32 and 79, 164 and 528, and 24 and 136 kg grain kg^−1^ nutrient for the SS zone. The minimum nutrient uptake coefficients (r) of N, P and K were 4.0, 0.5 and 4.5 kg ha^−1^ for the NGS zone, and 6.1, 0.8 and 7.3 kg ha^−1^ for the SS zone. In both agro-ecological zones, the parameterized QUEFTS model predicted a linear increase in above-ground dry matter uptake of N, P and K until the grain yield reached about 50-60% of the potential yield. At 60% of the potential yield (6 t ha^−1^) a balanced uptake in the above-ground part of 19.4 kg N, 3.3 kg P and 23.0 kg K are required to produce a tonne of maize grain in the NGS, and 17.7 kg N, 5.3 kg P and 26.2 kg K to produce a tonne of maize grain in the SS zone. Validation results indicated a good correlation between the parameterized QUEFTS estimated and observed grain yields in both agro-ecological zones. The parameterized QUEFTS is therefore a suitable model for balanced nutrient requirements estimation and development of site-specific fertilizer recommendations to improve maize yield in the Northern Nigerian Savanna. To ensure a greater impact, site-specific fertilizer recommendations must be complemented with appropriate agronomic management practices including use of the right source of fertilizer and a precise timing and placement of fertilizer application.

## Acknowledgements

The authors appreciate the financial support for this research from the Bill and Melinda Gates Foundation (BMGF) through the project called Taking Maize Agronomy to Scale in Africa “TAMASA” (contract ID: OPP1113374). We recognize the contribution of all field technologies of Centre for Drylands Agriculture (CDA), Bayero University Kano, Nigeria for coordinating the establishment, management and data collection of the trials. We also appreciate the efforts made by the laboratory staff of the International Institute for Tropical Agriculture (IITA), Nigeria for carrying out soil and plant analyses. All findings, conclusions and recommendations conveyed in this publication are those of the authors and do not necessarily reflect the view of the donor.

